# Muscle LIM Protein is a Non-Canonical RNA-Binding Protein that Engages in RNA-Dependent Cytoskeletal Assembly

**DOI:** 10.64898/2026.03.10.710776

**Authors:** Paul Sigrist, Dimitrios Destounis, Venkatraman Ravi, Sudeep Sahadevan, Thileepan Sekaran, Jonas Böhnlein, Verena Kamuf-Schenk, Zoe Loewenthal, Dunja Ferring-Appel, Frank Stein, Mandy Rettel, Etienne Boileau, Timon Seeger, Christoph Dieterich, Norbert Frey, Matthias W. Hentze, Mirko Völkers

## Abstract

Muscle LIM Protein (MLP) is a key component of the cardiomyocyte Z disc that is essential for sarcomere integrity and cardiac homeostasis. Accordingly, loss of MLP function results in (cardio)myopathy phenotypes in animal models and human patients.

In this study, we present the first RNA-binding proteome (RBPome) of human cardiomyocytes using enhanced RNA interactome capture (eRIC). Data integration with existing RBPome datasets identified MLP as an evolutionary conserved and previously unrecognized non-canonical RNA-binding protein (RBP) in cardiomyocytes. Using complementary biochemical approaches, we confirmed direct RNA association of MLP. Moreover, RNA integrity was required for MLP interactions with cytoskeletal proteins and for the formation of higher-order MLP-containing assemblies, revealed by co-immunoprecipitation and RNA-dependent sedimentation profiling. Subcellular fractionation and imaging analyses indicated that MLP’s association with cytoskeletal complexes may be weaker and more dynamic compared to core sarcomere proteins like α-actinin. Domain mapping identified two glycine-rich regions within MLP as RNA contact sites, and RNA-binding-deficient MLP mutants showed impaired association with cytoskeletal complexes. Together, our findings suggest RNA binding as a regulated property of MLP and uncover an RNA-dependent layer of cytoskeletal organization in cardiomyocytes.

## Introduction

RNA-binding proteins (RBPs) are essential regulators of cardiomyocyte function, and their dysregulation contributes to the development of cardiovascular disease. In recent years, systematic efforts to define the RNA-binding proteomes (RBPomes) of diverse tissues and cell types have revealed a surprisingly large cohort of RBPs whose capacity to directly engage RNA had previously gone unrecognized^1–3^. A shared characteristic of many of these non-canonical RBPs is the absence of well-defined evolutionary conserved RNA-binding domains (RBDs), suggesting that these proteins contact RNA via previously underappreciated mechanisms – typically with lower affinity and reduced specificity compared to canonical RBPs^4^. Notably, intrinsically disordered regions (IDRs) are strongly enriched within experimentally mapped RNA-binding regions of non-canonical RBPs^5^, supporting the emerging view that structural flexibility and low-complexity regions enable these proteins to interact with RNA in a context-dependent manner. Moreover, even established, canonical RBPs frequently harbor such non-canonical RNA-binding motifs adjacent to their canonical, structured RBDs, broadening their interaction potential and functional repertoire^6,7^.

In previous work, the RBPome of primary rat cardiomyocytes was identified, and critical regulatory roles of classical RBPs in controlling cell growth and survival were demonstrated^8–10^. However, functional studies of non-canonical RBPs in primary cells such as cardiomyocytes remain limited.

To elucidate how non-canonical RBPs control cardiomyocyte function, we took a multi-omics approach. We first obtained proteomic data about active RBPs in human iPS-derived cardiomyocytes (hiPS-CMs) and integrated those data with existing RBPome data from primary rat cardiomyocytes. This analysis identified Muscle LIM Protein (MLP; CSRP3) as a non-canonical RNA-binding protein in cardiomyocytes, with conservation observed across species.

MLP is a small, striated muscle-specific protein belonging to the cysteine-rich protein (CRP) family, which localizes to the sarcomeric Z disc. Genetic ablation of MLP in mice results in a phenotype reminiscent of human dilated cardiomyopathy (DCM), and mutations in the CSRP3 gene are linked to human cardiomyopathies highlighting its importance for long-term cardiac function^11,12^. Despite this clear physiological relevance, the molecular mechanisms through which MLP contributes to cardiomyocyte homeostasis remain incompletely understood, which motivated us to explore the functional consequences of RNA binding to MLP.

RNA-immunoprecipitation Sequencing (RIP-Seq) against MLP as well as proteomic analysis of the RNA-dependent MLP protein interactome in cardiomyocytes were combined and uncovered a previously unknown RNA-dependent mechanism of MLP targeting to the Z disc in cardiomyocytes. Those data suggested that MLP localization and interaction with Z disc proteins was dependent on its RNA binding, as an RNA-binding-deficient MLP mutant showed impaired association with cytoskeletal complexes. Thus, our data suggest a novel molecular mechanism of RNA-dependent assembly of sarcomeric protein complexes.

## Results

### Uncovering the human cardiomyocyte RBPome identifies MLP as an evolutionary conserved RBP

We previously reported the identification of novel cardiomyocyte-enriched RBPs in neonatal rat ventricular cardiomyocytes (NRVMs)^8^. To determine whether these RBPs are conserved in human cardiomyocytes and to prioritize non-canonical RBP candidates for functional characterization, we performed enhanced RNA interactome capture (eRIC) in human induced pluripotent stem cell–derived cardiomyocytes (hiPSC-CMs) (Fig. 1A). We subjected lysates from hiPS-CMs to eRIC in biological triplicates. eRIC eluates and the corresponding input samples were labeled by 10-plex tandem mass tag (TMT) and subjected to liquid chromatography-tandem mass spectrometry (LC-MS/MS). Analysis of proteins enriched in eluates after UV crosslinking compared to non-crosslinked control samples identified 341 proteins as RBP candidates, while more than 6000 proteins were detected in input samples, representing the total proteome (Fig. 1B). Among the 341 RBP candidates, we observed strong enrichment of proteins involved in RNA biology, as determined by gene ontology (GO) analysis (Fig. 1C). Integration with our previously established RBP atlas from NRVMs revealed conservation of 201 proteins between both datasets (Fig. 1D). These conserved RBPs as well showed significant enrichment for RNA-related GO terms (Fig. 1E). Interestingly, only two of the previous identified NRVM-specific RBPs were also identified as hits in hiPSC-CMs, suggesting that many of these may exhibit context-dependent RNA binding, potentially specific to certain species and maturation states (Fig. 1F). Notably, these two conserved non-canonical RBPs, MLP and Four and a Half LIM Domains Protein (FHL2), are closely related members of the LIM-only protein family with comparable reported biological roles as mechanosensitive cytoskeletal adaptor proteins. Furthermore, both are well-established disease genes associated with genetic cardiomyopathies^13^. Given the elevated expression of MLP observed in both human and murine heart failure, as well as the association between MLP mutations and genetic cardiomyopathies in human patients, we chose MLP for functional follow up experiments to investigate the functional implications of RNA binding to MLP.

**Fig. 1.**
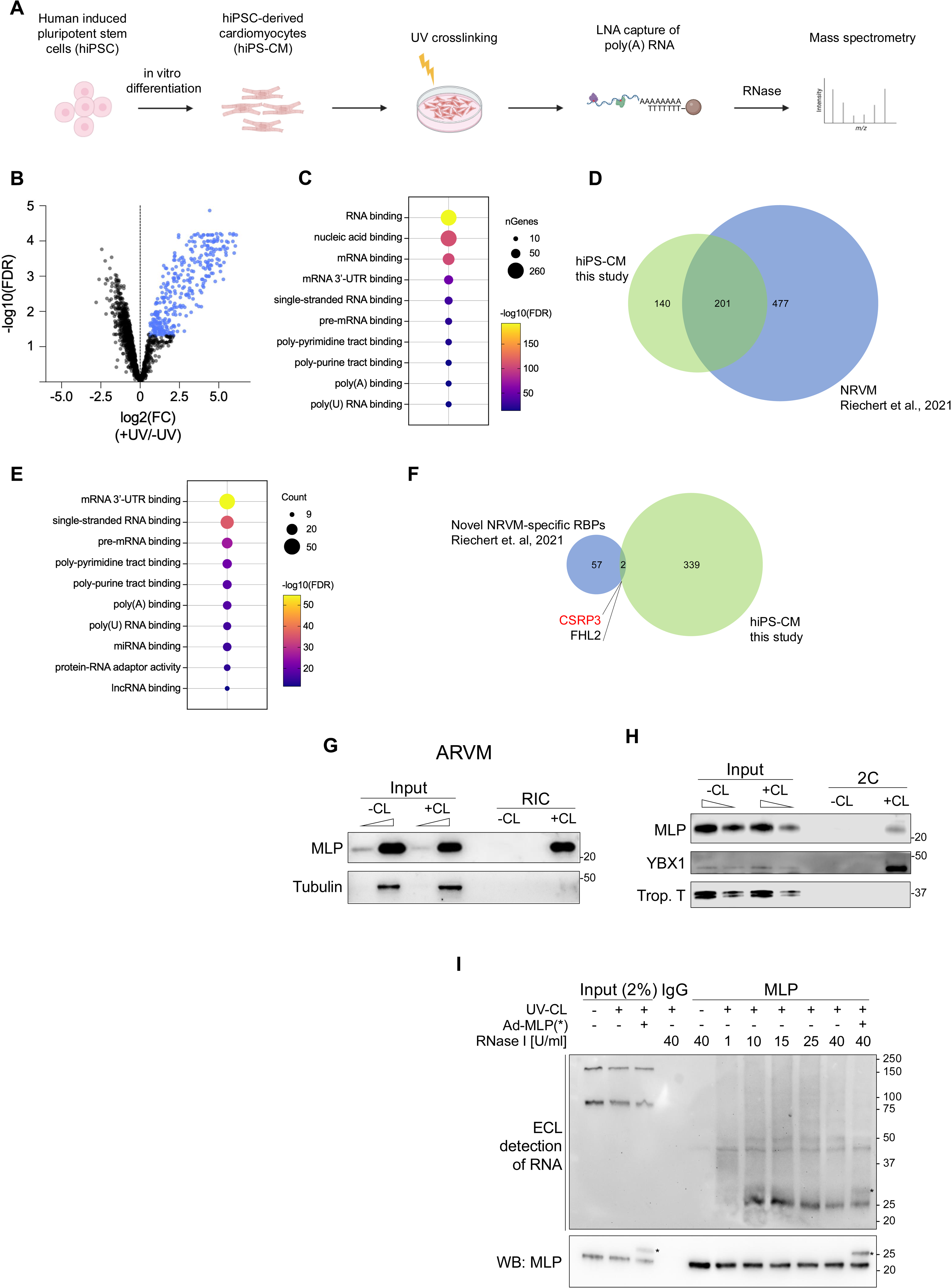
Uncovering the human cardiomyocyte RBPome identifies MLP as an evolutionary conserved RBP. (A) Workflow of eRIC in hiPS-CM. (B) Volcano plot of proteins detected in UV-crosslinked vs. non-crosslinked eluates. “Hit” RBPs (fold change >1.5, FDR <0.05) are marked in blue. (C) GO term analysis of enriched RBPs in hiPS-CM. (D) Venn diagram of overlap between RBPome datasets from NRVMs and hiPS-CMs. (E) GO term analysis of conserved RBPs. (F) Venn diagram of conservation of novel RBPs from NRVMs in hiPS-CMs. (G) Representative western blot analysis after RIC ARVM lysates. (H) Representative western blot after 2C in NRVMs. (I) Representative result of pCp assay for MLP in NRVMs.

Having identified MLP as an RBP candidate in NRVMs^8^ and hiPS-CMs through global proteomic analysis, we set out to obtain additional biochemical evidence for a direct interaction between MLP and RNA. We first validated the association of MLP with poly(A) RNA in both NRVMs and ARVMs in Western Blots after RIC (Fig. 1G). The orthogonal complex capture (2C) assay likewise recovered MLP from NRVM lysates, further suggesting direct UV-crosslinking (Fig. 1H). To independently confirm RNA binding, we applied a pCp assay to NRVM lysates – a non-radioactive adaptation of the established PNK assay, which relies on 3’-end ligation of a biotinylated nucleotide rather than 5’-end phosphorylation. This method consistently detected RNA in MLP immunoprecipitations (IPs) specifically after UV-crosslinking, with the MLP-RNA adducts migrating at the expected molecular weight (∼25 kDa) after extensive RNase I-mediated RNA trimming (Fig. 1I,J). Adenoviral overexpression of FLAG-tagged MLP resulted in a double band pattern in both western blot and streptavidin-mediated RNA detection, further supporting the specificity of the RNA signal.

### Identification of MLP target mRNAs *in vivo*

Global MLP knockout (KO) mice develop a dilated cardiomyopathy phenotype characterized by impaired systolic function and disrupted sarcomeric ultrastructure, previously establishing MLP as an essential regulator of cardiac mechanical integrity and stress response signaling *in vivo*^11^. To validate impaired cardiac function and structural remodeling in MLP KO mice compared to wild-type (WT) littermates under our conditions, we performed phenotypic analysis using echocardiography and histology. At 12 weeks of age, MLP KO mice showed a marked increase in left ventricular interior dimension, as shown in Masson’s trichrome-stained heart sections, and a significant decrease in systolic heart function assessed by transthoracic echocardiography (Fig. 2A-D). Heart weights were measured in hearts from KO versus WT mice and were unchanged (Fig. 2E), consistent with the previously reported dilated rather than hypertrophic phenotype. We measured the mRNA expression of marker genes for pathological cardiac remodeling by qRT-PCR and found increased levels of *Nppa* and *Nppb* as well as *MYH7* mRNA in MLP KO mice compared to WT (Fig. 2F).

**Fig. 2.**
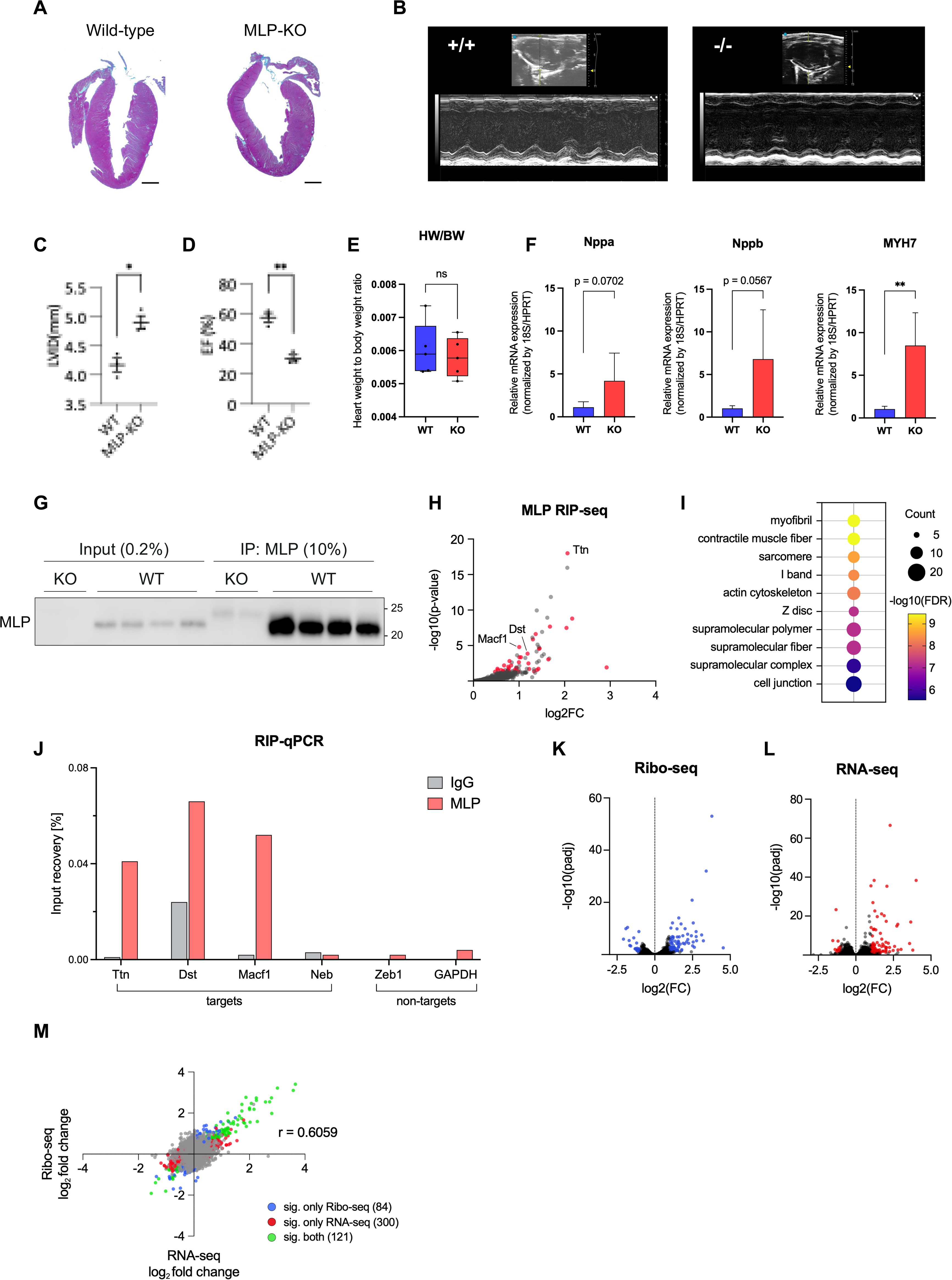
Identification of MLP target mRNAs *in vivo.* (A) Masson’s trichrome staining of WT and MLP KO hearts. Scale bars = 1 mm. (B) Echocardiography analysis of WT and MLP KO hearts. (C) Quantification of LVID from (B). n = 3 mice per group. *p < 0.05 by unpaired t test. (D) Quantification of ejection fractions (EF) from (B). n = 3 mice per group. **p < 0.01 by unpaired t test. (E) Heart weight to body weight ratios from WT and MLP KO mice. n = 5 mice per group. (F) qRT-PCR analysis of mRNA expression of *Nppa, Nppb* and *MYH7* in WT and MLP KO mice. n = 5 mice per group. **p < 0.01 by unpaired t test. (G) Western blot analysis confirming MLP capture in RIP-seq. (H) Enrichment volcano plot of enriched RNAs in MLP IPs from WT hearts over Input and IPs from KO hearts. Transcripts significantly enriched in WT IPs over WT input compared to KO IPs over KO input (log2FC > 0, p < 0.05) are highlighted in red. (J) RIP-qPCR validation of selected targets in NRVMs. n = 1. (K) Volcano plot Ribo-seq analysis of WT vs MLP KO mouse hearts. Significantly differently expressed genes (DEGs) (log2FC < -1 or > 1, p < 0.05) are highlighted in blue. (L) Volcano plot of Ribo-seq analysis of WT vs MLP KO mouse hearts. Significantly DEGs (log2FC < -1 or > 1, p < 0.05) are highlighted in red. (M) Scatter plot of integrated Ribo- and RNA-seq.

To systemically identify mRNAs associated with MLP, we performed native RNA-immunoprecipitation against MLP followed by high-throughput sequencing (RIP-Seq) of the immunopurified RNAs. RIP-Seq was performed from WT heart lysates as well as from MLP KO hearts which served as background controls. In parallel, RNA from all input samples was isolated and sequenced. RIP-Seq was performed in four biological replicates. Specific pull-down of MLP from WT hearts, but not from KO hearts, was validated by Western blot analysis (Fig. 2G).

RIP-seq analysis revealed 38 transcripts significantly enriched in MLP IPs from WT hearts over input and control IPs. Among these, we found significant enrichment of mRNAs encoding for sarcomeric and structural proteins in cardiomyocytes, including *Titin* (*Ttn*) and *Desmin* (*Des*) (Fig. 2H,I). Binding of MLP to selected candidates was validated by conducting native RIP for endogenous MLP in NRVMs. Quantification of co-purified transcripts was assessed by qRT-PCR. Indeed, RIP-qPCR confirmed enrichment of target mRNAs detected in RIP-seq compared to non-target transcripts in MLP IPs, but not in IPs using a species-matched IgG isotype control antibody.

We further aimed to determine putative regulatory effects of MLP binding on its target mRNAs as well as to study translational changes induced by loss of MLP on a global scale. To this end, we sequenced mRNAs occupied by actively translating ribosomes using Ribo-Seq from WT and MLP KO hearts (Fig. 2K)^14^. Only periodic fragment lengths that showed a distinctive triplet periodicity were kept for downstream analysis. In parallel, input samples from the same hearts were subjected to RNA-seq for transcriptomic analysis (Fig. 2L).

In total, we identified 505 differentially expressed genes by combined Ribo- and RNA-seq in MLP KO hearts. Integration of both datasets revealed that MLP KO did not result in significant changes in global translational efficiency, as changes in translation correlated strongly with changes in mRNA expression (Fig. 2M).

### Cytoskeletal interactions of MLP are RNA-dependent

Since regulatory directionality in non-canonical RBPs is not necessarily centered on the control of RNA fate, and because we did not observe clear regulatory dominance of MLP over the translation of its mRNA targets *in vivo*, we aimed to investigate whether RNA binding might instead modulate MLP function. To date, a global characterization of MLP’s protein interactome has been lacking, and only a limited number of specific protein-protein interactions (PPIs) have been mapped^15,16^. Therefore, our goal was to generate a comprehensive dataset defining MLP’s interaction network and to identify interactions that might be regulated by RNA. To this end, we compared native MLP Co-IPs from untreated NRVM lysates with Co-IPs performed on lysates pretreated with RNase I to deplete RNA prior to IP (Fig. 3A).

**Fig. 3.**
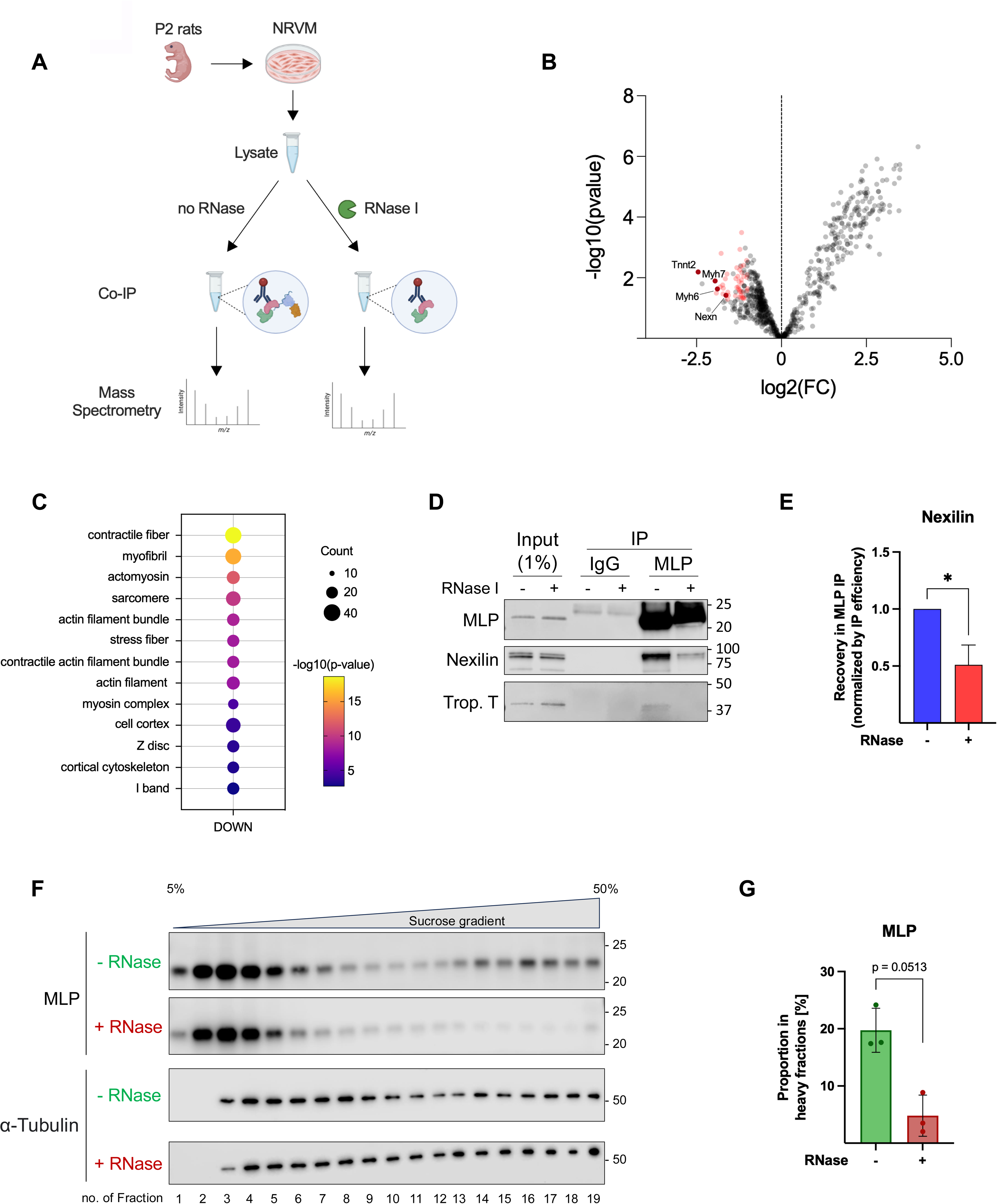
Cytoskeletal interactions of MLP are RNA-dependent. (A) Workflow and mechanistic rationale of MLP Co-IP with RNase digestion. (B) Volcano plot of proteins in MLP Co-IP after RNase vs untreated comparison. Proteins that show a significant decrease in MLP IP upon RNase treatment compared to untreated samples (log2FC < -1; p < 0.05) are highlighted in red. (C) GO term analysis of proteins significantly reduced in MLP IP after RNase vs untreated IPs. (D) Representative western blot of selected RNase-sensitive interactors. (E) Quantification of Nexilin recovery in MLP IPs dependent on RNase treatment. n = 3. *p < 0.05 by paired t test. (F) R-DeeP sucrose gradient centrifugation of NRVM lysates followed by western blot for MLP.

Surprisingly, we observed that a substantial number of MLP interactions with sarcomeric proteins were sensitive to RNase, displaying markedly reduced co-recovery with MLP following RNA digestion (Fig. 3B). GO-term analysis of proteins differentially recovered between RNase-treated and untreated conditions further supported this observation, indicating that predominantly interactions with cytoskeletal components were lost upon RNA removal (Fig. 3C). We validated a set of representative candidates including canonical sarcomere proteins like Nexilin and Troponin T by Western Blotting (Fig. 3D-E). Conversely, we detected increased recovery of multiple RNA-associated proteins, including mostly ribosomal proteins, possibly reflecting their enhanced availability for unspecific MLP binding in cell lysates due to RNase-mediated disassembly of endogenous RNP complexes.

To independently assess MLP’s participation in RNA-dependent higher-order assemblies, we performed R-DeeP analysis on NRVM lysates (Fig. 3F,G)^17^. Comparison of sucrose gradient profiles from untreated samples and samples subjected to RNase I digestion prior to ultracentrifugation revealed an RNase-mediated redistribution of a distinct subpopulation of MLP. Under basal conditions, this fraction sedimented in heavier sucrose fractions and consistently shifted toward lighter fractions upon RNA degradation. In contrast, the majority of MLP localized to low-density fractions irrespective of RNase treatment, indicating that RNA-dependent assembly may involve only a subset of the total MLP pool, consistent with lower stoichiometry of MLP’s RNA association compared to canonical RBPs^18^.

### MLP dynamically associates with the actin cytoskeleton

Given that several cytoskeletal interactions of MLP were RNase-sensitive, we next examined its association with the actin cytoskeleton and sarcomeres by immunostainings. In sections of adult rat hearts, MLP immunostaining resulted in a cross-striated pattern colocalized with the canonical Z disc protein α-actinin (Fig. 4A,B). Consistent with previous reports in NRVMs^19^, endogenous MLP displayed a predominantly diffuse staining pattern (Fig. 4C), in contrast to the highly ordered Z disc localization observed in adult cardiomyocytes and adult cardiac tissue sections. In addition to this diffuse signal, we detected filamentous MLP structures that partially overlapped with actin filaments visualized by phalloidin staining (Fig. 4C). We hypothesized that the diffuse signal may represent a soluble cytoplasmic pool of MLP. To better resolve cytoskeleton-associated MLP, NRVMs were subjected to a classical protocol for subcellular fractionation prior to fixation employing a buffer designed for cytoskeletal preparations containing 0.5% Triton X-100. Interestingly, this treatment eliminated the majority of the MLP signal, while preserving actin filaments (Fig. 4C). Extraction of MLP protein into the soluble fraction together with cytoplasmic proteins and retention of canonical sarcomere proteins including α-actinin, cardiac myosin binding protein C (Mybpc3) and Troponin T in the insoluble fraction were further validated by western blotting (Fig. 4D). This observed increased sensitivity of MLP to detergent-extraction compared to core sarcomeric components is in line with previous reports in both NRVMs and adult cardiac tissue^19^ and supports a rather weak, transient and potentially RNA-mediated association of MLP with cytoskeletal complexes. Together with the prominent diffuse signal observed in intact cells, these findings indicate that MLP associates with the cytoskeleton in a comparatively weak and dynamic manner, in line with its suspected role as a load-responsive signaling component rather than a core structural element.

**Fig. 4.**
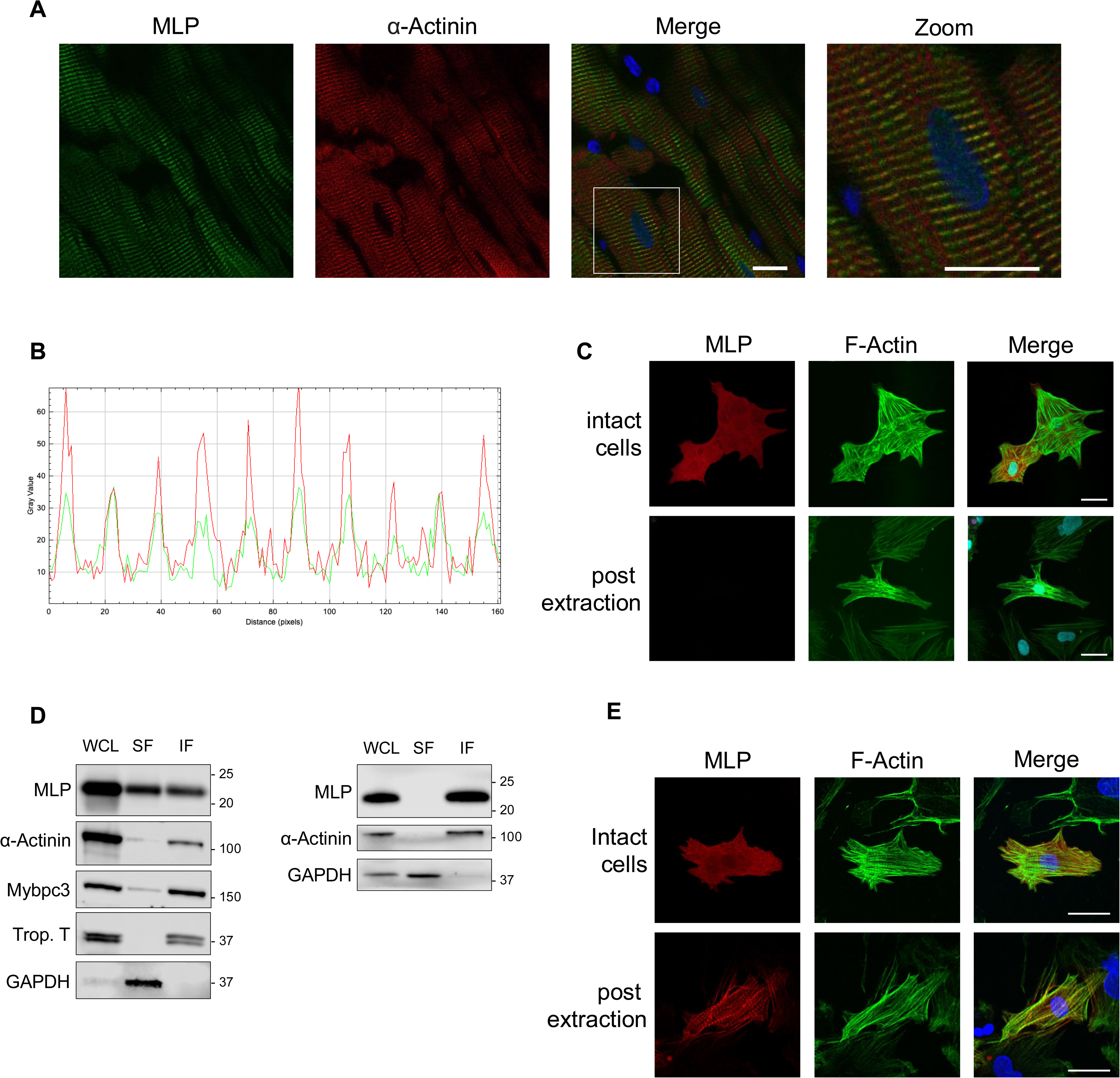
MLP dynamically associates with the actin cytoskeleton. (A) Immunofluorescence staining of MLP and α-Actinin in adult rat heart sections (Scale bar = 25 µm). (B) Line scan analysis of composite image of MLP (green) and α-Actinin (red) from (A). (C) Representative western blot analysis after subcellular fractionation of NRVMs. (left) Fractionation using a buffer containing 0.5% Triton X-100 as detergent. (right) Fractionation using Buffer I of the ProteoExtract subcellular proteome extraction kit for extraction of cytosolic proteins. (D) Representative immunofluorescence images of MLP and F-actin (stained by phalloidin-fluorescein) in NRVMs before (top row) and after (bottom row) extraction using the Triton X-100 buffer. (E) Representative immunofluorescence images of MLP and F-actin in NRVMs before (top row) and after (bottom row) cytosol extraction using the ProteoExtract subcellular proteome extraction kit.

When testing different extraction protocols to preserve cytoskeletal MLP, we found that cytosolic extraction using the ProteoExtract subcellular proteome extraction kit preserved cytoskeletal MLP, while extracting cytosolic proteins like GAPDH (Fig. 4D). Interestingly, extraction of the soluble protein pool eliminated the diffuse signal observed in intact cells and instead unmasked cytoskeletal MLP. While in many cells the MLP signal still appeared predominantly filamentous and co-localized with phalloidin stained actin filaments, many cells also showed cross-striated MLP signal reminiscent of the pattern observed in adult tissue sections, which was not detectable in intact cells (Fig. 4E). However, since western blot analysis of the soluble (SF) and insoluble (IF) fractions did not show extraction of a detectable proportion of MLP protein (Fig. 4D), our data do not support a model in which a large pool of MLP is cytoplasmic and freely soluble. Instead, we speculate that removal of soluble factors may influence binding equilibria of dynamic interactions and may thereby force MLP into a cytoskeleton-bound state, again supporting a highly transient interaction mode.

### MLP contacts RNA via two glycine rich regions

Since MLP lacks canonical RBDs, we next aimed to identify protein regions mediating its interaction with RNA. MLP contains two structured LIM domains, each followed by flexible glycine-rich regions (GRRs). Published RBDmap data indicate that the MLP paralog CSRP1 contains a potential RNA-interacting region near the boundary of the LIM2 domain and the C-terminal glycine-rich region (GRR2), inferred from peptides detected near RNA-crosslinked residues (Npeps)^20^. To generate independent data for MLP, we performed a domain-mapping approach using a series of truncation mutants of MLP fused to a C-terminal 3×FLAG tag (Fig. 5B). RNA-binding capacity was assessed by comparative pCp assay analysis following adenoviral expression of the different constructs with specific deletions in differentiating C2C12 myotubes (Fig. 5C). Notably, RNA–protein crosslinked species consistently migrated around 15 kDa above the expected molecular weight in pCp assays following anti-FLAG-tag IP compared to conventional western blotting; while the precise origin of this shift remains unclear, its strict dependence on construct size and domain composition clearly supports its specificity.

**Fig. 5.**
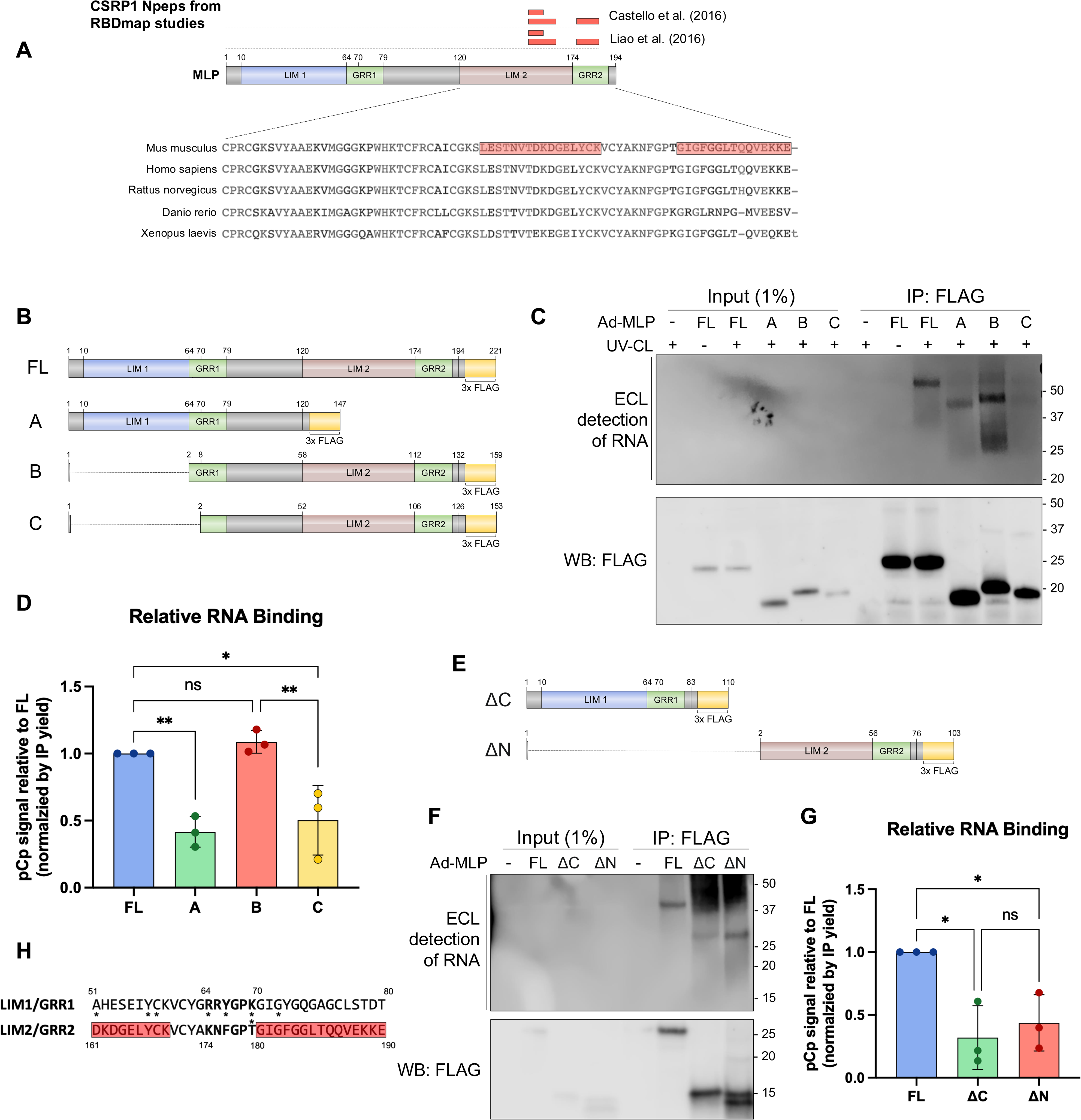
MLP contacts RNA via two glycine rich regions. (A) Scheme of MLP’s domain architecture and mapping of published RBDmap data for CSRP1 on the MLP sequence. (B) Scheme of FLAG-tagged MLP constructs used for domain mapping. (C) Representative pCp assay after adenoviral expression of domain constructs in differentiating C2C12 myotubes. (D) Quantification of RNA binding to domain constructs assessed by pCp assays in C2C12. N = 3. *p < 0.05, **p < 0.01 by one-way ANOVA. (E) Scheme of N- and C-terminal MLP constructs. (F) Representative pCp assay after adenoviral expression of ΔC-MLP and ΔN-MLP in NRVMs. (G) Quantification of RNA binding to ΔC-MLP and ΔN-MLP assessed by pCp assays in NRVMs. N = 3. *p < 0.05 by one-way ANOVA. (H) Sequence comparison of LIM/GRR1 and LIM2/GRR2 boundaries. Bold letters mark experimentally determined residues crucial for RNA binding in the N-terminal module (aa 64-69) and the analogous stretch in the C-terminal module (aa 174-179). Red boxes represent Npeps identified for CSRP1. Asterisks indicate sites of mutations reported in HCM or DCM patients.

Deletion of the N-terminal 10 amino acids together with the LIM1 domain (construct B) did not significantly affect RNA association. In contrast, removal of the C-terminal 74 amino acids encompassing the LIM2 domain (construct A), as well as deletion of a short six–amino-acid stretch (aa 64–69) immediately downstream of the LIM1 domain (construct C), resulted in a marked reduction in RNA crosslinking (Fig. 5C,D). This latter region has previously been described as a functional nuclear localization signal (NLS)^21^. These findings suggested that at least two distinct regions contribute to RNA binding and that each region alone can mediate RNA interaction, albeit with reduced efficiency compared to full-length MLP.

To further test this hypothesis, we generated additional constructs comprising either LIM domain together with its adjacent GRR. pCp assay analysis following expression in NRVMs confirmed that both the N-terminal (ΔC) and C-terminal (ΔN) modules are independently capable of RNA binding, with comparable apparent affinity (Fig. 5E-G), but with lower binding capacity compared to full-length MLP.

Given the apparent structural redundancy, we next focused on identifying shared sequence features that might underlie RNA affinity. Integration of our mapping data with the published RBDmap data revealed that the Npeps identified for CSRP1 flank a short sequence motif containing a six–amino-acid stretch analogous to the region downstream of LIM1 identified here (Fig. 5H), suggesting that these analogous regions may represent MLP’s two independent RNA binding motifs.

### Impaired cytoskeletal targeting of an RNA binding deficient MLP mutant

To directly test the contribution of these regions to RNA binding, we generated a mutant of the N-terminal module by exchanging the six amino acids we identified to be crucial for RNA binding (aa 64-69). We paid specific attention to remove aromatic and basic (positively charged) residues, which are crucial mediators in canonical nucleic acid binding domains, and replaced those with hydrophobic and negatively charged residues (Fig. 6A). Following adenoviral expression in NRVMs, RNA binding was assessed by pCp assays. In support of our proposed model, mutation of these six amino acids resulted in a marked reduction in RNA-binding (Fig.6B,C) to MLP. We next examined the subcellular behavior of the RNA-binding–deficient mutants in NRVMs. Biochemical fractionation revealed a pronounced change in fractionation behavior. While FLAG-tagged wt-MLP was found to remain mostly insoluble, similar to endogenous MLP, the mutant protein was almost completely solubilized under these conditions suggestive of a reduced or weakened sarcomeric association (Fig. 6D-E). Since this finding could potentially be explained by disruption of a specific PPI through the introduced mutations, we tested the fractionation behavior of the single module constructs (ΔC and ΔN). Both truncated proteins showed increased solubility comparable to the RNA binding deficient mutant (Fig. 6F).

**Fig. 6.**
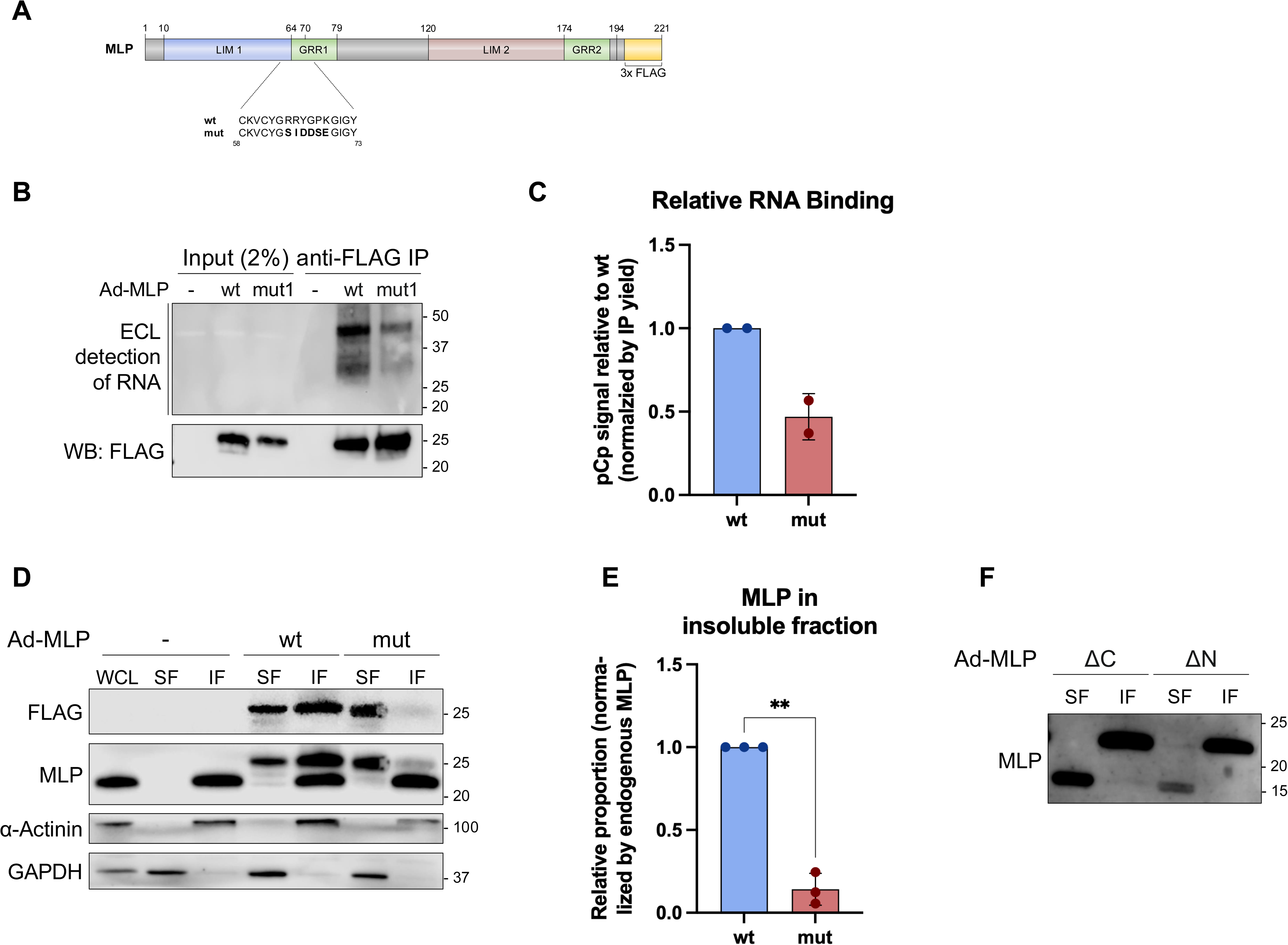
Impaired cytoskeletal targeting of an RNA binding deficient MLP mutant. (A) Scheme of mutagenesis strategy. (B) Representative pCp assay after adenoviral expression of wt and mutant MLP in NRVMs. (C) Quantification of RNA binding assessed by pCp assays. N = 2. (D) Representative western blot after subcellular fractionation following adenoviral expression of wt and mutant MLP in NRVMs. (E) Quantification of D. n = 3. *p < 0.05 by paired t test. (F) Western blot of fractionation after expression of ΔC-MLP and ΔN-MLP in NRVMs.

Collectively, these data indicate that efficient integration of MLP into its sarcomeric interaction network in primary cardiomyocytes is likely not mediated by a single PPI but instead relies on a complex and potentially multivalent binding mode which may involve RNA binding mediated by two short flexible regions.

## Discussion

Muscle LIM protein, encoded by the *CSRP3 gene*, is a crucial component of the sarcomeric Z disc in cardiomyocytes, and has previously been assumed to function mainly as a scaffold protein that assembles macromolecular complexes and signaling complexes driving cardiac remodeling ^9,14,17^. It has been further suggested that MLP may act as a mechanosensor at the Z disc, where mechanical stress may trigger its nuclear translocation, potentially linking altered cardiac load to changes in gene expression and cardiac remodeling^17^.

Mutations in the CSRP3 gene are thought to disrupt MLP’s essential, but incompletely understood functions, leading to myofibrillar disorganization, fibrosis, and severe cardiac disease such as hypertrophic (HCM) and dilated cardiomyopathy (DCM) in human patients. In addition, global deletion of the MLP gene in mice results in a severe DCM phenotype. Consistent with a conserved role in muscle biology, homologs of MLP have been shown to be essential for normal muscle development and function in other organisms, including Drosophila and zebrafish, where loss of function results in comparable phenotypes, underscoring its evolutionary importance^23,24^. However, how these conserved functions are mechanistically executed at the molecular level remains unclear.

In this study we identify MLP as an evolutionary conserved non-canonical RBP and present evidence supporting an RNA-mediated mechanism that governs MLP’s subcellular localization and its incorporation into the Z disc network. Molecular mapping indicates that MLP’s RNA binding ability is mediated by short flexible regions adjacent to the structured LIM domains. Notably, the corresponding region within the LIM1/GRR1 module overlaps with a cluster of reported pathogenic MLP variants in humans (asterisks in Fig. 5H), suggesting that this region may be functionally important. Of note, impaired RNA binding is unlikely to represent the main molecular mechanism for all disease-associated mutations. For example, the C58G mutation identified in HCM patients affects one of the zinc ion-coordinating cysteine residues in the N-terminal LIM1 domain of MLP and has been reported to destabilize the protein through misfolding and increased degradation^25^.

A long-standing controversy concerns the subcellular distribution and functional mode of action of MLP. While numerous studies have described MLP as a structural Z disc protein in cardiomyocytes, others have reported a predominantly cytoplasmic localization^19^ and in some studies nuclear accumulation has been observed^21^. In skeletal muscle progenitors, MLP has been proposed to act as a co-factor for transcription factors (TFs) during myoblast differentiation, suggesting a potential nuclear function^26^, which has not been demonstrated in cardiomyocytes. Collectively, these observations have led to multiple, sometimes conflicting models of MLP function, ranging from structural support to signal modulation and transcriptional regulation.

Mechanistically, the two adjacent glycine-rich regions immediately next to the LIM domains are crucial for RNA binding to MLP. LIM domains have recently been proposed as a previously overlooked class of RBDs exclusively based on the detection of numerous LIM domain–containing proteins in several RNA interactome studies^27^. While our data support RNA binding of LIM domain–containing proteins such as MLP, our mapping analyses strongly indicate a LIM domain–independent mechanism of RNA interaction. This observation suggests that the recurrent identification of LIM domain proteins in RBPome datasets may rather reflect the frequent coexistence of LIM domains with (adjacent) independent RNA-binding motifs than direct RNA binding mediated by the LIM domains themselves. A comparable principle has been described for TFs of the homeodomain family, in which DNA binding is mediated by the homeodomain while associated LIM domains primarily contribute to TF activity by shaping the protein interaction network. In conclusion, our data in primary cardiomyocytes indicate that the presence of each LIM domain alone is insufficient to promote stable association with sarcomeric structures, further supporting a model in which RNA binding - rather than a single PPI - critically contributes to MLP’s cytoskeletal engagement in a physiological context.

We identified several mRNAs encoding for structural and sarcomeric proteins binding to MLP. Integrated RNA-seq and Ribo-seq analyses indicated that deletion of MLP does not have a significant impact on the translational efficiency of its target mRNAs. These results further support the notion that MLP binding to mRNA might primarily play a role in intracellular localization rather than translation regulation.

In summary, we identify RNA-binding as a new function of MLP in cardiomyocytes. MLP-associated mRNAs include transcripts encoding huge and key structural proteins like titin. As many of these transcripts are thought to undergo localized translation at the sarcomeric Z disc^28^, it is very tempting to speculate that such long and also long-lived RNAs might – besides their role as templates for sarcomere protein synthesis – exert non-canonical functions including scaffolding cytoskeleton-associated protein complexes within the densely crowded Z disc network. While the exact molecular mechanism how RNA contributes to Z disc targeting of MLP remains elusive, the finding that several other Z disc and sarcomere proteins have been detected in RBPome studies suggests that RNA-dependent assembly may represent a more general concept with potential implication in Z disc organization, calling for further investigation. Our findings offer a direct molecular connection of RNA-binding to MLP localization to the Z disc. MLP binding to mRNA might, therefore, be essential for proper sarcomeric structure and myofibrillar organization. Further research is needed for a better understanding of how differential RNA binding behavior of MLP is regulated and how MLP binding to mRNA at the Z disc is regulated in response to neurohumoral stimulation. Moreover, validating the functional impact of RNA binding *in vivo* using relevant mouse models remains crucial and requires further investigation to better understand RNA interactions with non-canonical RPBs and their effects on pathological remodeling of cardiomyocytes.

## Methods and Materials

### Cell culture

Neonatal rat ventricular cardiomyocytes (NRVMs) were isolated by enzymatic digestion of 1–4 d-old neonatal rat hearts and purified by Percoll density gradient centrifugation before plating, as previously described^14^. Tissue culture plates were pre-coated with 0.1% gelatine for 1 h at 37°C. Briefly, cardiac myocytes were plated in DMEM/F-12 medium containing 10% fetal bovine serum (FBS). After 24 h media were replaced with DMEM/F12

Adult rat Ventricular myocytes (ARVMs) from adult rat hearts were isolated using a collagenase method. Animals were anesthetized with and the aorta was rapidly cannulated after the hearts were excised and perfused with a rate of 8 ml/min in a Langendorff apparatus. Hearts were initially perfused with calcium-free medium (pH 7.2) consisting of (in mM) 5.4 KCl, 3.5 MgSO4, 0.05 pyruvate, 20 NaHCO3, 11 glucose, 20 HEPES, 23.5 glutamate, 4.87 acetate, 10 EDTA, 0.5 phenol red, 15 butanedionemonoxime (BDM), 20 creatinine, 15 creatine phosphate (CrP), 15 taurine and 27 units/ml insulin under continuous equilibrium with 95% O2/5% CO2. After 5 min collagenase was added (0.5 U/ml, Worthington) for 20 – 30 min. Left ventricles of digested hearts were cut into small pieces and subjected to gentle agitation to allow for dissociation of cells. Consequently, cells were resuspended in M199 without BDM in which 2 mM extracellular calcium was gradually reintroduced.

Human iPSCs were cultured in feeder-free and serum-free conditions in DMEM/F12 medium (Corning, 15-090-CM) supplemented with homemade E8 supplement on Matrigel-coated plates (growth factor-reduced, Corning, 354230) in a humidified incubator at 37 °C and 5% CO2. Homemade E8 supplement was prepared by combining: L-ascorbic acid 2-phosphate (64 mg/l, Sigma-Aldrich, A92902-25G), Insulin (19.4 mg/l, Life Technologies, 17504-001), transferrin (10.7 mg/l, Sigma-Aldrich, T8158-100MG), sodium selenite (14 µg/l, Sigma-Aldrich, S5261-10G), FGF2 (100 µg/l, Life Technologies, 100-18B-50UG) and TGFβ1 (2 ng/l, Life Technologies, 100-21-10UG). Differentiation of human iPSC into CMs was performed using a small-molecule Wnt-activation/inhibition protocol^29^. For the first 7 days, human iPSCs were differentiated using cardiac differentiation medium (CDM3) composed of RPMI1640 (Life Technologies, 21875091) supplemented with L-ascorbic acid 2-phosphate (0.213 g/l, Sigma-Aldrich, A92902-25G) and recombinant human albumin (0.5 g/l, Sigma-Aldrich, A9511-100MG) in sterile water. From day 1-3, the iPSCs were cultured in CDM3 medium containing a gradient concentration of CHIR99021 (4-2 µmol/l, TOCRIS, 4953). From day 3-5, media was changed to CDM3 medium supplemented with IWP2 (5 µmol/l, Med Chem Express, 686770-61-6), followed by CDM3 only from day 5-7. From day 7 onward, cells were cultured in RPMI1640 supplemented with B27 (Life Technologies, 17504044). From day 10-13, hiPSC-CM were maintained in RPMI1640 without glucose, supplemented with CDM3 supplement and lactate. At day 13-15, beating human iPSC-CM were either cryopreserved using Cryo-Brew freezing medium (Miltenyi Biotec, 130-109-558), or passaged using TrypLE select enzyme (Life Technologies, A1217702) followed by resuspension in passaging medium RPMI1640 containing B27, knockout serum replacement (Life Technologies, 10828028) and Rock inhibitor (Sigma-Aldrich, Y0503-1MG), and reseeding on Matrigel-coated plates. eRIC experiments were conducted using human iPSC-CM with >90% purity between day 45 and 50 after the initiation of differentiation.

C2C12 myoblasts were cultured in DMEM media. For pCp assays, fresh vials of cells were thawed and seeded in DMEM media supplemented with 15% FBS. 24h after plating, the cells were split and plated on 150 mm plastic dishes in DMEM media containing 15% FBS. To induce differentiation, media were replaced by differentiation media (DMEM containing 2% horse serum). After one day of differentiation, cells were transduced with the respective adenovirus in differentiation media. Media were changed daily and cells were harvested after 3 days of differentiation.

### Enhanced RNA-Interactome Capture (eRIC)

eRIC was performed as previously described^30^. hiPS-CMs were UV-crosslinked and lysed in RIC Lysis Buffer (20 mM Tris-HCl pH 7.5, 500 mM LiCl, 1 mM EDTA, 5 mM DTT, 0.5% (w/v) LiDS). Lysates were homogenized by rigorous needling, snap frozen in liquid nitrogen and stored at -80°C until use. 18.75 x 10^6^ hiPS-CMs were used per replicate.

### Complex capture (2C)

Cells were washed twice with ice-cold PBS and UV-crosslinked (150 mJ/cm2 for cell lines and 240 mJ/cm2 for primary cells) on ice before lysis. Non-irradiated samples were used as controls. Lysates were sonicated using a Bioruptor (5-10 cycles, 30 sec on, 30 sec off, mode: high) and centrifuged for 10 min at 14.000 rpm at 4°C to pellet cell debris. The protein concentration of the resulting supernatants was quantified and the samples were stored at - 80°C until use.

Complex capture was carried out using the Zymo© Quick-RNA Miniprep Kit (#R1055) reagents according to the manufacturer’s recommendations. Two successive rounds of column capture were performed. In between both rounds of 2C^2^ the eluates of the first column were treated with 10 U TURBO™ DNase (#AM2238) for 30 min at 37°C in a 100 µl reaction. Where stated 500 U of Ambion™ RNase I (#AM2295) were added to this reaction for some samples to serve as an additional RNA-depleted control.

Following the second round of purification RNA-protein complexes were eluted from the columns, treated with 10.000 U/ml of RNase I (500 U in 50 µl) for 30 min at 37°C and prepared for Western Blotting.

### pCp-Assay

Cell lysates were prepared as described for 2C^2^. For IP, Dynabeads Protein G (#10004D; for mouse antibodies) or Dynabeads Protein A (#10002D; for rabbit antibodies) were washed twice in iCLIP Lysis Buffer and incubated with primary antibody (20 µl of bead slurry and 4 µg of antibody for 1 mg of lysate) over night at 4°C. Samples were thawed and RNA was partially digested using mild (1-40 U/ml) RNase I digestion (15 min, 37°C, 850 rpm). After RNase digestion 440 U/ml (11 µl per ml) of RNase Inhibitor (RNase Inhibitor, Murine (NEB), #M0314L) were added and the samples were again centrifuged (12.000 rpm, 10 min, 4°C). After antibody-bead coupling the beads were washed thrice in iCLIP Lysis Buffer and incubated with the lysates for 2h at 4°C to capture protein complexes. After IP beads were washed once with iCLIP Lysis Buffer, two times with iCLIP High Salt (HS) Wash Buffer, once with iCLIP LiCl HS Wash Buffer and twice with iCLIP Wash Buffer. During the high salt washed the beads were put on a rotator for 2-3 min per wash. Transition washes were performed from iCLIP Lysis Buffer to iCLIP HS Wash Buffer and from iCLIP LiCl HS Wash Buffer to iCLIP Wash Buffer. Then beads were washed once in FAST AP Wash Buffer before starting on bead phosphorylation in a two-step incubation:

1. Beads were treated with Alkaline Phosphatase (FAST AP; #EF0651) and TURBO™ DNase for 30 min at 37°C, 850 rpm.
2. T4 Polynucleotide Kinase (T4 PNK; #M0201L) was added together with 5x PNK Buffer pH 6.5 and beads were incubated further for 20 min at 37°C, 1.000 rpm interval shaking (for 15 sec every two min).

After dephosphorylation RNA fragments covalently crosslinked to IPed proteins were biotin labeled using T4 RNA Ligase 1 (ssRNA ligase, High Concentration; #M0437M) and pCp-Biotin (#NU-1706-BIO) as substrate. Beads were incubated in labeling solution for 2h at 16°C.

For elution beads were finally resuspended in 21 µl 0.2 M Glycine pH 2.0 for 10 min at 1.200 rpm at RT. Beads were magnetically separated and the eluates were pH neutralized with 4 µl 1 M Tris-Cl pH 8.0.

Samples were denatured by adding 4x Laemmli Buffer (#1610747) plus 100 mM. RNA-Protein complexes were resolved on an SDS-PAGE and transferred to a nitrocellulose membrane. Biotinylated RNA-adducts on proteins were visualized using the Chemiluminescent Nucleic Acid Detection Module Kit (#89880).

### Co-immunoprecipitation

Cells were lysed in Co-IP Lysis Buffer (20 mM Tris-Cl pH 7.5, 150 mM NaCl, 2 mM MgCl_2_, 10% Glycerol, 0.5% NP40) and homogenized by passing the lysates through needles of decreasing diameters. The lysates were centrifuged, and the supernatants were aliquoted at a protein concentration of 1 mg/ml. For Co-IP experiments Dynabeads were washed twice in Co-IP Lysis Buffer and incubated with primary antibody over night at 4°C. 20 µl of bead slurry and 4 µg of antibody were used per 1 mg of lysate. Isotype control IgG IPs served as controls.

Following antibody-bead coupling the beads were washed thrice in Co-IP Lysis Buffer and incubated with the lysates for 2h at 4°C to capture protein complexes. After IP, beads were washed three times with Co-IP Lysis Buffer by inverting the tubes. For elution, beads were finally resuspended in 21 µl 0.2 M Glycine pH 2.0 for 10 min at 1.200 rpm at RT. Beads were magnetically separated and the eluates were neutralized with 4 µl 1 M Tris-Cl pH 8.0.

To test RNA-dependence of PPIs, lysates were digested with 500 U/ml RNase I for 30 min at 37°C immediately before starting the IP. Untreated control samples were also incubated for 30 min at 37°C.

### Mass spectrometry data acquisition and analysis

Protein samples were subjected to the SP3 protocol^31^ conducted on the KingFisher Apex™ platform (Thermo Fisher). For digestion, trypsin was used in a 1:20 ratio (protease:protein) in 50 mM N-2-hydroxyethylpiperazine-N-2-ethanesulfonic acid (HEPES) supplemented with 5 mM Tris(2-carboxyethyl)phosphine hydrochloride (TCEP) and 20 mM 2-chloroacetamide (CAA). Digestion was carried out for 5 hours at 37°C.

Up to 10 µg of peptides were labeled using TMTpro™ 16plex reagent as previously described^32^. Briefly, 0.5 mg of TMT reagent was dissolved in 45 µL of 100% acetonitrile. Subsequently, 4 µL of this solution was added to each peptide sample, followed by incubation at room temperature for 1 hour. The labeling reaction was quenched by adding 4 µL of a 5% aqueous hydroxylamine solution and incubating for an additional 15 minutes at room temperature. Labeled samples were then combined for multiplexing, desalted using an Oasis® HLB µElution Plate (Waters) according to the manufacturer’s instructions, and dried by vacuum centrifugation.

Offline high-pH reversed-phase fractionation^33^ (PMID: 22462785) was carried out using an Agilent 1200 Infinity high-performance liquid chromatography (HPLC) system, equipped with a Gemini C18 analytical column (3 μm particle size, 110 Å pore size, dimensions 100 x 1.0 mm, Phenomenex) and a Gemini C18 SecurityGuard pre-column cartridge (4 x 2.0 mm, Phenomenex). The mobile phases consisted of 20 mM ammonium formate adjusted to pH 10.0 (Buffer A) and 100% acetonitrile (Buffer B). The peptides were separated at a flow rate of 0.1 mL/min using the following linear gradient: 100% Buffer A for 2 minutes, ramping to 35% Buffer B over 59 minutes, increasing rapidly to 85% Buffer B within 1 minute, and holding at 85% Buffer B for an additional 15 minutes. Subsequently, the column was returned to 100% Buffer A and re-equilibrated for 13 minutes. During the LC separation, 48 fractions were collected. These were pooled into six fractions by combining every sixth fraction. The pooled fractions were then dried using vacuum centrifugation.

An UltiMate 3000 RSLCnano LC system (Thermo Fisher Scientific) equipped with a trapping cartridge (µ-Precolumn C18 PepMap™ 100, 300 µm i.d. × 5 mm, 5 µm particle size, 100 Å pore size; Thermo Fisher Scientific) and an analytical column (nanoEase™ M/Z HSS T3, 75 µm i.d. × 250 mm, 1.8 µm particle size, 100 Å pore size; Waters) was used. Samples were trapped at a constant flow rate of 30 µL/min using 0.05% trifluoroacetic acid (TFA) in water for 6 minutes. After switching in-line with the analytical column, which was pre-equilibrated with solvent A (3% dimethyl sulfoxide [DMSO], 0.1% formic acid in water), the peptides were eluted at a constant flow rate of 0.3 µL/min using a gradient of increasing solvent B concentration (3% DMSO, 0.1% formic acid in acetonitrile). Peptides were introduced into an Orbitrap Fusion™ Lumos™ Tribrid™ mass spectrometer (Thermo Fisher Scientific) via a Pico-Tip emitter (360 µm OD × 20Lµm ID; 10 µm tip, CoAnn Technologies) using an applied spray voltage of 2.2LkV. The capillary temperature was maintained at 275 °C. Full MS scans were acquired in profile mode over an m/z range of 375–1,500, with a resolution of 120,000 at m/z 200 in the Orbitrap. The maximum injection time was set to 50 ms, and the AGC target limit was set to ‘standard’. The instrument was operated in data-dependent acquisition (DDA) mode, with MS/MS scans acquired in the Orbitrap at a resolution of 30,000. The maximum injection time was set to 94 ms, with an AGC target of 200%. Fragmentation was performed using higher-energy collisional dissociation (HCD) with a normalized collision energy of 34%, and MS2 spectra were acquired in profile mode. The quadrupole isolation window was set to 0.7 m/z, and dynamic exclusion was enabled with a duration of 60 seconds. Only precursor ions with charge states 2–7 were selected for fragmentation.

Raw files were converted to mzML format using MSConvert from ProteoWizard, using peak picking, 64-bit encoding and zlib compression, and filtering for the 1000 most intense peaks. Files were then searched using MSFragger in FragPipe (22.1-build02) against FASTA database UP000002494_RattusNorvegicus_BrownNorway_ID10116_22816entries_26102022_dl1101202 3 containing common contaminants and reversed sequences. The following modifications were included into the search parameters: Carbamidomethylation (C, 57.0215), TMTpro (K, 304.2072) as fixed modifications; Oxidation (M, 15.9949), Acetylation (protein N-terminus, 42.0106), TMTpro (peptide N-terminus, 304.2072) as variable modifications. For the full scan (MS1) a mass error tolerance of 20 PPM and for MS/MS (MS2) spectra of 20 PPM was set. For protein digestion, ’trypsin’ was used as protease with an allowance of maximum 2 missed cleavages requiring a minimum peptide length of 7 amino acids. The false discovery rate on peptide and protein level was set to 0.01. The standard settings of the FragPipe workflow ’Default’ were used. For the proteomics data analysis the raw output files of FragPipe (protein.tsv files files) were processed using the R programming environment (ISBN 3-900051-07-0). Initial data processing included filtering out contaminants and reverse proteins. Only proteins quantified with at least 2 razor peptides (with Razor.Peptides >= 2) were considered for further analysis. 723 proteins passed the quality control filters. Log2 transformed raw TMT reporter ion intensities values (’channel’ columns) were normalized using the ’normalizeVSN’ function of the limma package^34^ . Normalization was performed separately for the following groups: IgG and MLP. This ensured that abundance differences across these groups were preserved. Missing values were imputed with the ’knn’ method using the ’impute’ function from the Msnbase package^35^. This method estimates missing data points based on similarity to neighboring data points, ensuring that incomplete data did not distort the analysis.

For the proteomics data analysis the raw output files of FragPipe (protein.tsv files files) were processed using the R programming environment (ISBN 3-900051-07-0). Initial data processing included filtering out contaminants and reverse proteins. Only proteins quantified with at least 2 razor peptides (with Razor.Peptides >= 2) were considered for further analysis. 6826 proteins passed the quality control filters. In order to correct for technical variability, batch effects were removed using the ’removeBatchEffect’ function of the limma package^34^ on the log2 transformed raw TMT reporter ion intensities (’channel’ columns). Subsequently, normalization was performed using the ’normalizeVSN’ function of the limma package (VSN - variance stabilization normalization)^36^. Missing values were imputed with the ’knn’ method using the ’impute’ function from the Msnbase package^35^. This method estimates missing data points based on similarity to neighboring data points, ensuring that incomplete data did not distort the analysis. Differential expression analysis was performed using the moderated t-test provided by the limma package. The model accounted for replicate information by including it as a factor in the design matrix passed to the ’lmFit’ function. Imputed values were assigned a weight of 0.01 in the model, while quantified values were given a weight of 1, ensuring that the statistical analysis reflected the uncertainty in imputed data. To obtain p-values and false discovery rates (FDRs), the ’fdrtool’ function from the fdrtool package^37^ was used to analyze the t-values produced by limma for certain comparisons. Proteins were annotated as hits if they had a false discovery rate (FDR) below 0.05 and an absolute fold change greater than 1.5. Proteins were considered candidates if they had an FDR below 0.2 and an absolute fold change greater than 1.5. Clustering with all enriched hit proteins based on the median protein abundances normalized by median of control condition was conducted to identify groups of proteinsimilar patterns across conditions. The ’kmeans’ method was employed, using Euclidean distance as the distance metric and ’ward.D2’ linkage for hierarchical clustering. The optimal number of clusters was determined using the Elbow method, which identifies the point where the within-group sum of squares stabilizes. Gene ontology (GO) enrichment analysis was performed using the ’compareCluster’ function of the ’clusterProfiler’ package^38^, which assesses over-representation of GO terms in the dataset relative to the background gene set. The analysis was performed using ’org.Hs.eg.db’ as the reference database. The odds ratio (’odds_ratio’) for each GO term was calculated by comparing the proportion of genes associated with that term in the dataset (’GeneRatio’) to the proportion in the background set (’BgRatio’). An odds ratio greater than 1 indicates that the GO term is enriched in the dataset compared to the expected background.

### Immunoblotting

Samples were combined with the appropriately concentrated form of Laemmli sample buffer and then boiled before SDS-PAGE followed by transfer to PVDF or nitrocellulose membranes. The membranes were probed with the following antibodies: CSRP3/MLP (1:1000, XXX), YBX1 (1.1000, Cell signaling technology, #sc-398340), TnnT (1:000), GAPDH (1:5000, Santa Cruz sc-365062),

### RNA-immunoprecipitation

Left ventricular cardiac tissue was harvested from wt and MLP KO mice and snap frozen in liquid nitrogen. Frozen tissue was then lysed in RIP Lysis Buffer (20 mM Tris-HCl pH 7.5,150 mM KCl, 2 mM MgCl_2_ 0,5% NP-40) and triturated by passing through needles of decreasing diameters (19 – 27 G) for complete lysis. CSRP3 antibody (Proteintech, cat. no. 10721-1-AP) was coupled to Dynabeads Protein A (Invitrogen, cat. no. 10002D) over night. Lysates were then incubated with prepared beads for 2 h at 4°C for RNP capture. Following IP, beads were washed three times with RIP Lysis Buffer. RNA was isolated using the RNeasy mini kit (Quiagen, cat. no. 74104) according to the manufacturer’s instructions after eluting the beads in 200 µl of RLT buffer. Takara SMART-Seq Total RNA Library Prep was used for library generation following the manufacturer’s instructions. Libraries were multiplexed and sequenced on a NextSeq550.

### RIP-Seq analysis

Reads were trimmed with Cutadapt (v2.5) and mapped to rat genome (Rnor_6.0) with STAR (v2.7) and summarized with featureCounts(v1.6.4). DESeq2 (Love et al., 2014) with IHW (Ignatiadis et al., 2016) for multiple hypothesis correction was used to determine significantly enriched RNAs in IP samples vs corresponding input controls (adjusted p-value < 0.05; log2 fold-change > 0.5).

### Polysome profiling and RiboSeq

Ribo-seq libraries were prepared for five biological replicates as published previously (Doroudgar et al., 2019). Ribosome footprints were generated after isolation of polyribosomes from LV lysates and RNase I digestion as previously published (Kmietczyk et al., 2019). Briefly, left ventricular tissues of WT and MLP KO mice were lysed in polysome buffer (20 mM Tris pH 7.4, 10 mM MgCl, 200 mM KCl, 2 mM DTT, 100 μg/ml CHX, 1% Triton X-100, 1U DNAse/μl) containing 100 μg/ml CHX. For complete lysis, the samples were kept on ice for 10 min and subsequently centrifuged at 20,000 × g to pellet cell debris and the supernatant was immediately used for further preparation. Sucrose solutions were prepared in polysome gradient buffer and 20 U/mL SUPERase-In. Sucrose density gradients (10–50% wt/vol) were freshly made in SW40 ultracentrifuge tubes using a BioComp Gradient Master (BioComp). Cell lysates were loaded onto sucrose gradients, followed by centrifugation at 220,000 × g for 250 min at 4 °C in an SW40 rotor. Separated samples were fractionated at 0.375 ml/min by using a fractionation system BioComp Gradient Station (BioComp) that continually monitors OD254 values. Monosomal fractions were collected into tubes at 0.3 mm intervals. Libraries were generated according to the mammalian NEXTFLEX Small RNA-Seq Kit v4 (Revvity) and sequenced on a NextSeq 550.

### RiboSeq-Analysis

We followed our previously published protocols (Doroudgar et al.,2019). Ribo-seq data was analyzed with Ribotools (https://ribotools.readthedocs.io). Abundance was estimated from ribosome-protected fragments using periodic reads only. UMI, A-tail, template switch motif and adapters sequences were trimmed using Flexbar with "-u 2 -q TAIL -qt 10 -qf sanger --htrim-right AAAAAA --pre-trim-left 16 -k 248 -m 15 -g -d -ao 6" according to the D-plex Small RNA-seq Kit guidelines. For the statistical analyses, we used the edgeR package. We only considered data points with read count observations across all replicates. We used an FDR<0.05 and FC=2 (log2 FC=1) as cutoff.

### qRT-PCR

Quantitative RT-PCR was performed in duplicate measurements on samples from input and RIC eluate RNA and amplified using specific oligonucleotide primers designed to the indicated transcripts primers.

For quantification of 18S rRNA and gapdh mRNAs reverse transcription (RT) followed by RT-qPCR was performed. RT was performed using the iScript™ cDNA Synthesis Kit (Bio-Rad). cDNA from total noCL RNA was used as standard. Amplification was performed in the presence of 1LmM of each oligonucleotide and iTaq Universal SYBR Green Supermix (Bio-Rad). Relative amounts of targets were calculated using the ddCT method. Sequences of oligonucleotides used for quantification of 18S and gapdh mRNA were designed with Primer3 software.

### Immunofluorescence and staining

Immunostaining of myocytes was performed as described before (Volkers et al., 2013). NRVMs were grown on gelatin-coated permanox chamber slides or 14 mm glass coverslips. Cells were fixed by 4% paraformaldehyde (PFA) for 10 min at room temperature, washed three times with PBS, and permeabilized in PBS with 0.3% Triton-X for 10 min, then blocked in PBS with 10% horse serum for 1h. Primary antibodies diluted in blocking solution (PBS with 10% horse serum) were applied overnight at 4 °C. The next day, cells were washed with PBS and incubated for 1h at RT with secondary antibody (Jackson Laboratories) diluted in blocking solution. After three washing steps in PBS, cells were mounted in Vectashield containing 1:1000 DAPI as nuclear staining.

### Immunocytofluorescence of adult rat heart sections

Paraffin-embedded hearts were sectioned and placed on slides, which were then deparaffinized and rehydrated. Antigen retrieval was achieved by boiling the slides in 10 mM citrate (pH 6.0) for 12 min, after which the slides were washed several times with distilled water, and once with Tris/NaCl, or TN buffer (100 mM Tris and 150 mM NaCl). Primary antibodies were diluted in TNB and added to slides which were incubated at 4°C for ∼12–16 h. The samples were then washed with TN buffer and incubated with secondary antibodies at room temperature in the dark for 2 h. Masson Trichrome stainings were performed as per manufacturer instructions (Trichomes Accustain (Masson), Sigma-Aldrich). Images were obtained using a Zeiss Observer.Z1 fluorescence microscope. Images were obtained with a 63× objective.

### Statistics

Cell culture experiments were performed at least two times with n = at least two biological replicates (cultures) for each treatment. In vivo experiments were performed on at least three biological replicates (mice) for each treatment. Cell size measurements of RVCMs or heart sections were performed with at least two biological replicates and at least 50 cells measured per replicate. The investigators have been blinded to the sample group allocation during the experiment and analysis of the experimental outcome. Unless otherwise stated, values shown are mean ± SEM and statistical treatments are one-way ANOVA followed by Bonferroni’s post hoc comparisons (three or more experimental groups) or unpaired t-test (two experimental groups) as indicated in the figure legends.

## Acknowledgments

We thank all members of the MV. and MWH laboratory for helpful discussion and comments. P.S., D.D., C.D, N.F. and M.V. acknowledge the DZHK (German Centre for Cardiovascular Research) Partner Site Heidelberg/Mannheim. M.V. is supported by the Heisenberg Programm of the Deutsche Forschungsgemeinschaft. P.S., D.D, T.S. M.V., M.W.H. and N.F. were supported by Collaborative Research Center 1550 (CRC1550/SFB1550) “Molecular Circuits of Heart Disease.”

## Author Contributions

P.S., M.W.H. and M.V. conceptualized the project, designed experiments, analyzed data, and wrote the manuscript. S.S., T.S., E.B, J.B., T.S, C.D. and N.F. analyzed data and provided critical input. P.S., D.D, V.R., V.K.S., Z.L., M.R. and F.S. performed experiments.

## Declaration of Interests

The authors declare no competing interests.

## Notes

### Competing Interest Statement

The authors have declared no competing interest.

